# Lab-Based and Self-Reported Indices of Fitness Show Lowered Fitness and Insight into Fitness in Individuals at Clinical High Risk for Psychosis

**DOI:** 10.1101/2020.04.30.060244

**Authors:** Katherine S. F. Damme, Richard P. Sloan, Matthew N. Bartels, Alara Ozsan, Luz H. Ospina, David Kimhy, Vijay A. Mittal

**Author notes:** **Corresponding Author:** Katherine Damme, Department of Psychology, Northwestern University. Denotes equal contribution of authorship (David Kimhy, Vijay A. Mittal).

## Abstract

**Introduction:** Exercise is a promising intervention for clinical high-risk for psychosis (CHR) populations, who have attenuated positive symptoms, but evidence suggests that these youth may require tailored exercise interventions. Presently, the scope of the problem is unknown, as these youth may not be reliable reporters on fitness. This issue is compounded by the fact that there have been no investigations that utilized a formal fitness assessment in this critical population. The present study aims to determine the level of fitness in CHR youth with lab-based measures, test how effectively self-report measures characterize objective fitness indices, and explore clinical factors that may be interrupting reliable self-report-an important tool if these interventions are to be taken to scale.

**Methods:** Forty CHR individuals completed an exercise survey and lab-based indices of fitness (i.e., VO_2_max and BMI). Forty healthy volunteers completed lab indices of fitness and a structured clinical interview ruling out the presence of psychiatric illness.

**Results:** CHR youth showed greater BMI and lower VO_2_max compared to healthy volunteers. In the CHR group, abstract self-report items (perceived fitness) did not reflect lab indices of fitness, whereas specific exercise behaviors (intensity of exercise) showed stronger correlations with laboratory-based fitness measurements. Exploratory analyses suggested that positive symptoms involving grandiosity, and negative symptoms such as avolition, correlated with discrepancy between self-perception and laboratory findings of fitness.

**Discussion:** Results suggest that CHR individuals are less objectively fit than matched controls, and that it will be important to consider unique population characteristics when weighing self-report data.

## Introduction

Exercise is a promising new area of intervention for individuals across many stages of psychosis, including for those at clinical high risk for psychosis (CHR).^1–8^ Despite the benefits of this new area of intervention, substantially less work has examined unique factors of the CHR population that would have bearing on tailoring an effective, early intervention (i.e., understanding characteristic health and starting fitness levels).^7,9–11^ Extant research, however, has relied solely on self-report measures of physical activity to assess fitness,^9,10,12–14^ with few exceptions.^9,14^ However, such reports are vulnerable to the influence of episodic memory difficulties, emotional states at the time of the assessment, as well as cognitive biases/reframing.^15,16^ This issue is particularly acute in studies of individuals with schizophrenia, as well as at risk for psychosis, given the substantial episodic memory deficits documented in these populations.^17,18^ Additionally, symptoms of distortions of reality (i.e., grandiosity) may impact perceived fitness along with deficits in motivation (i.e., avolition). To address these gaps in the literature, the current study will examine multiple lab-based metrics of fitness, as well as self-reported measures of fitness in individuals at CHR for psychosis. Collectively, this information can serve to refine both assessments of fitness and targeted fitness intervention.

A growing body of research has demonstrated that exercise interventions are effective in improving cognitive function^1,4,5,11,19^ and neurological function^2,11^ for individuals across the psychosis spectrum including, risk mental states, CHR, the first episode of psychosis, and schizophrenia populations. Although existing psychosis literature has established that individuals with psychosis have significantly poorer physical health,^1,6,14,19,20^ the evidence regarding the physiological health of individuals at CHR for psychosis is unclear.^21,22^ Despite some evidence that CHR individuals are objectively less active^9^ than their peers, a number of studies suggest that this does not translate into lower levels of biometric fitness, i.e., a normal range of body mass index (BMI).^23–26^ However, it is critical to note that the existing examinations of BMI have been demographic descriptions for exclusion purposes;^23–26^ as a result BMI is not well characterized in CHR individuals. In contrast, CHR individuals consistently self-report lower levels of fitness, less physical activity, and more barriers to exercise.^10–14^ Clarifying the nature and extent of fitness deficits in early risk for psychosis would provide critical insight to developing and refining targeting of exercise interventions in these populations.^6,7,22,27^

These inconsistencies may be driven by methodological issues surrounding assessments of fitness. Alternatively, it is possible that biometric evaluations of fitness do not capture the most relevant features of fitness for CHR, and that physiological measures of fitness, such as VO_2_max, may be more sensitive assessments of emerging health issues.^21,22^ Although only one study to date has examined physiological fitness (i.e., VO_2_max) in individuals identified as offspring of individuals with psychosis, individuals with psychotic-like experiences, and/or future conversion to psychosis, this work found that VO_2_max was significantly lowered in these individuals compared to peers.^14^ This difference in physiological fitness among other risk populations^14^ may indicate that physiological health metrics are more sensitive assessments for risk populations such as CHR.^20,22,27^ Finally, it is possible that inconsistencies in fitness are driven by issues related to a self-report approach; self-report accounts of fitness behaviors have been found to have limited validity in capturing actual health behaviors^28–30^ which may be exacerbated by psychosis-like experiences,^31^ such as distortions in self-perception and memory. In the CHR individuals, several symptoms have good face validity in terms of being candidates for confounding accurate recall and reporting. In particular, symptoms such as grandiosity may serve to influence perceptions of physical fitness and experiences of avolition may distort experiences and reports around physical effort.

The current study is the first to examine if CHR show reduced objective, physiological and biometric markers of fitness compared to healthy volunteers. This study also investigates the validity of self-report measures in reflecting the actual fitness of CHR individuals by comparing lab-based indices of fitness with self-reported levels of perceived fitness and fitness behaviors. These analyses provide critical insight into whether objective markers of fitness provide added clinical insight beyond self-report measures of physical fitness. Additionally, analyses will also investigate whether attenuated psychotic symptoms impact self-reported experiences of fitness and perception of fitness. Finally, follow up analyses will explore whether symptom severity contributes to the discrepancies between self-report and lab-based metrics of physical fitness. The degree of mismatch between physical fitness and actual fitness (participant error) may reflect specific symptoms that impact the subjective experience of physical activity due to) to distortions of self (i.e., grandiosity) or the impairments in motivation (i.e., avolition). This relationship may inform exercise interventions for CHR individuals in how best to track performance and how perception may not reflect the reality of fitness in individuals at CHR.

## Methods

### Participants

The current study sample were combined from independent studies of aerobic fitness. The CHR individuals were collected at The Adolescent Development and Preventive Treatment (ADAPT) lab and the healthy volunteer sample were selected as a best match to that sample from a larger study conducted at the New York State Psychiatric Institute (NYSPI) at the Columbia University Medical Center (CUMC). All study procedures were conducted in accordance with the Northwestern University and NYSPI's Institutional Review Board approved guidelines, respectively, and all participants provided written informed consent. In the ADAPT lab, 40 CHR individuals completed a structured clinical interview assessing attenuated psychotic symptoms for inclusion into the study and to rule out the presence of a psychotic disorder. CHR subjects also completed an exercise survey (e.g., current exercise practices, perceived physical fitness, and lab-based indices of fitness, i.e. VO_2_max and BMI). At Columbia University, 40 gender and age matched healthy volunteers completed lab indices of fitness and a structured clinical interview ruling out the presence of psychiatric illness. Participants were recruited from advertising (healthy volunteers); inclusion criteria required healthy volunteers to meet American College of Sports Medicine's standards for exercise^32^ to ensure safety during participation in the fitness assessment.

### Clinical Assessment of Symptoms

The Structured Clinical Interview for DSM-IV Axis I Disorders (SCID) were administered to all subjects to rule out any psychosis diagnosis for both the CHR and healthy volunteer groups. CHR participants completed the Structured Interview for Psychosis Risk Syndromes (SIPRS) in order to assess the presence of a CHR syndrome, and to track the presence of attenuated symptoms. This study conducted a series of exploratory analyses on positive and negative domains, as well as two specific symptoms including grandiosity (an exaggerated sense of superiority/belief in special powers) and avolition (impaired motivation).

### Self-Reported Fitness Scale

CHR participants completed a self-report survey comprised of items from a number of validated measures^33–36^ and subscales of this self-report have been previously reported in assessing physical activity in CHR individuals.^10,12^ The items selected for the current study include: perceived fitness, frequency of exercise, time spent exercising, and intensity of exercise. Perceived fitness was rated by participants on a Likert-type scale that ranged from 0 (Poor) to 3 (Excellent). Frequency of exercise, where exercise was defined as any activity that resulted in sweating or rapid heartrate, was rated based on a typical week and ranged from 0 (rarely or never) to 3 (five or more times). Time spent exercising was rated on a scale spanning 0 (less than 30 minutes) to 3 (more than 60 minutes). Intensity of exercise assessed the frequency that these exercise sessions resulted in sweating or rapid heartrate on a scale from 0 (never) to 3 (always; every time).

### Lab-based Indices of Aerobic and Biometric Fitness

Lab based indices of fitness included VO_2_max and Body Mass Index (BMI). VO_2_max indexes an individual’s ability to transport and use oxygen at the maximum capacity during aerobic exercise. At Northwestern, an expert exercise physiologist conducted a modified Balke max-exercise protocol^37^ under the supervision of a physician. In this modified Balke max-exercise protocol, the treadmill speed was set to elicit 70% of age-predicted max heart rate and an RPE rating of around 13 (“somewhat hard”). During the protocol the speed of the treadmill remained the same, but the incline of the treadmill belt increased 2% every 2 minutes (or 2.5% for speeds 6 mph or greater). Tests generally lasted 8-12 minutes, the recommended target for VO_2_max testing.^38^ At Columbia, the Human Performance Laboratory at Columbia University Medical Center assessed the VO_2_max of participants during cycling. To generate the VO_2_max (mL/kg/min) variable, participants achieved one of the following four criteria during exercise: VO_2_ plateau, 85% of maximal heart rate (220-age), respiratory quotient ≥ 1.1, or self-reported exhaustion as indexed by the Borg Scale^39^ to generate the VO_2_max (mL/kg/min). For more detail on the full exercise procedure, refer to Kimhy et al.^1^ To ensure comparability across sites and exercise methods, gender and age appropriate z-scores were created for all VO_2_max scores according to the American Heart Association^40^ national norms. At both sites, body mass index (BMI) was calculated based on measurements of height and weight collected in lab using the U.S. Department of Health and Human Services’ National Heart Lung and Blood Institute (http://nhlbisupport.com/bmi/bminojs.html) BMI online calculator.^41^ Error of Fitness Estimation was calculated by relating lab-based fitness metrics to self-reported perceived fitness. The standard model error was computed for each subject, reflecting the difference between the actual values of self-reported perceived fitness and the expected values of fitness according to lab-based fitness metrics. Individuals were then grouped categorized into inaccurate and accurate reporter groups consistent with their quartile distribution of standardized errors. Those who were inaccurate in their estimates (either highly over- or under-estimating their fitness) were grouped as inaccurate reporters; those with more typical amounts of error in their estimates (near the 50^th^ percentile of standard model error) were grouped as accurate reporters for follow up analyses.

### Statistical Approach

Participant demographic comparisons across groups were conducted on sample features using chi-squared analyses to characterize categorical sample features and t-tests to examine continuous features of the sample. Any significant differences between groups will be used as nuisance covariates in subsequent analyses. Group comparisons (i.e., CHR and healthy volunteers) of lab-based metrics of health were conducted in separate simultaneous general linear models, where group will be used to predict fitness (VO_2_max or BMI, respectively) while accounting for variance related to demographics of the study sample. Analyses within the CHR group examined whether self-report indices of fitness related to lab-based indices; to limit the total number of analyses, self-report subscales were entered simultaneously into separate general linear models to predict VO_2_max and BMI in separate analyses. This approach has the added benefit of accounting for multiple self-reported features of exercise. As a result, any significant subscale will be significant while accounting for other features of self-reported fitness behaviors, providing added insight into the relevance of particular items over and above other items. A similar approach was taken regarding the impact of clinical features on self-reported indices of fitness; to reduce the number of total comparisons, a repeated-measure general linear model examined the relationship between clinical symptoms (positive and negative) as the within-subject measure and simultaneously the subscales of self-reported indices of fitness were entered as the between-subjects measure in a single model. In this approach a significant relationship to symptoms in general would indicate that the self-reported indices of fitness related to symptom severity, and an interaction by scale would indicate that the relationship varies by symptom type (positive or negative). Finally, in a set of exploratory analyses, CHR individuals were grouped by error quintiles as accurate or inaccurate (based on relationship of their perceived fitness to each lab-based fitness) and were compared to symptoms of grandiosity and avolition in separate t-tests: grandiosity by VO_2_max accuracy quintile, grandiosity by BMI accuracy quintile, avolition by VO_2_max accuracy quintile, avolition by BMI accuracy quintile. In significant analyses, follow-up analyses will be conducted to examine if sex or age contribute significantly to the models and demographics will be included; if the variable significantly contribute to the model or impacts the direction or magnitude of the reported findings.

## Results

### Participants

Our sample included 80 participants (healthy volunteers=40, CHR=40), *Table 1*. There were not significant differences in sex by group, χ^2^=0.46, *p*=.49. Although the groups used distinct exercise methods (i.e., treadmill, cycling), the peak heart rate was not significantly different, *t*(78)=0.131, *p*=.89, across CHR (*M*=181.61 *StD*=17.93) and healthy volunteers (*M*=181.13 *StD*=11.73); suggesting the exercise was of comparable rigor. There was a significant difference in age between groups, *t*(78)=−9.19, *p*<.001, where CHR individuals were younger (*M*=20.95, *StD*=1.38) than healthy volunteers (*M*=23.98, *StD*=1.56). Accordingly, all group comparisons will account for age related differences.

**Table 1.**
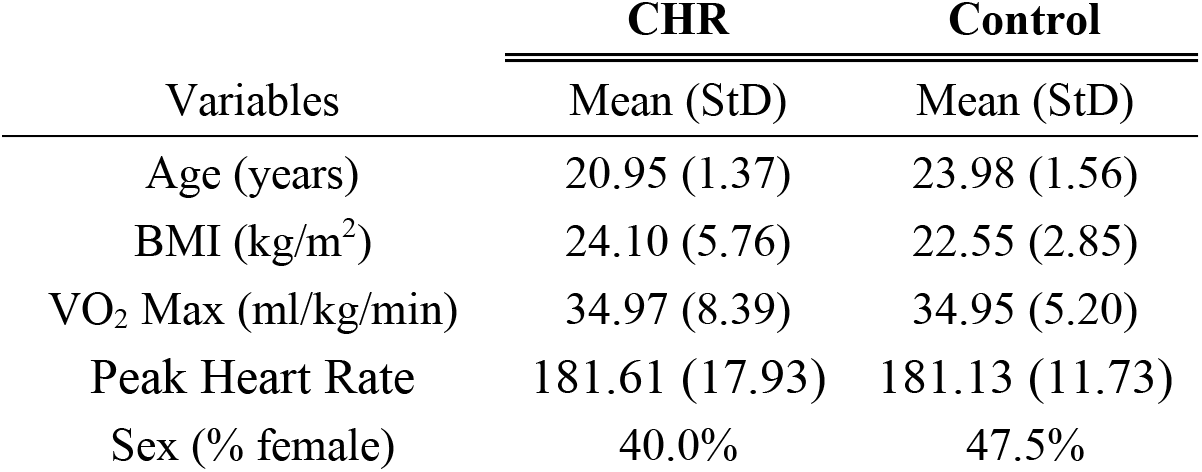
Group Demographics and Lab Based Exercise Metrics

| Variables | CHR | Control |
| --- | --- | --- |
|  | Mean (StD) | Mean (StD) |
| Age (years) | 20.95 (1.37) | 23.98 (1.56) |
| BMI (kg/m<sup>2</sup>) | 24.10 (5.76) | 22.55 (2.85) |
| VO<sub>2</sub> Max (ml/kg/min) | 34.97 (8.39) | 34.95 (5.20) |
| Peak Heart Rate | 181.61 (17.93) | 181.13 (11.73) |
| Sex (% female) | 40.0% | 47.5% |

### Group Differences Lab-Based Indices of Fitness

In a general linear model, group (CHR vs Healthy Volunteer) related to normed VO_2_max (ml/kg/min) z-scores, accounting for variability related to age. There was a significant main effect of group, *F*(1,77)=4.01, *p*=.049, such that the CHR group (*M*=−.97, *StD*=.178) had significantly lower VO_2_max compared to healthy volunteers (*M*=−.38, *StD*=.178), indicating reduced physiological health, *Figure 1a*. There was also a significant main effect of age, *F*(1,77)=5.99, *p*=.017, such that older age was related to lower VO_2_max values, *r_-partial_*=−.27. In a general linear model, group (CHR vs Healthy Volunteers) related to BMI (kg/m^2^), accounting for variability related to age. There was a significant main effect of group, *F*(1,77)=12.29, *p*=.001, such that the CHR group (*M*=25.75, *StD*=.84) had significantly increased BMI compared to healthy volunteers (*M*=20.91, *StD*=.84), indicating reduced health, *Figure 1b*. There was also a significant main effect of age, *F*(1,77)=12.29, *p*=.001, such that older age was related to increased BMI values, *r_-partial_*=.35.

**Figure 1.**
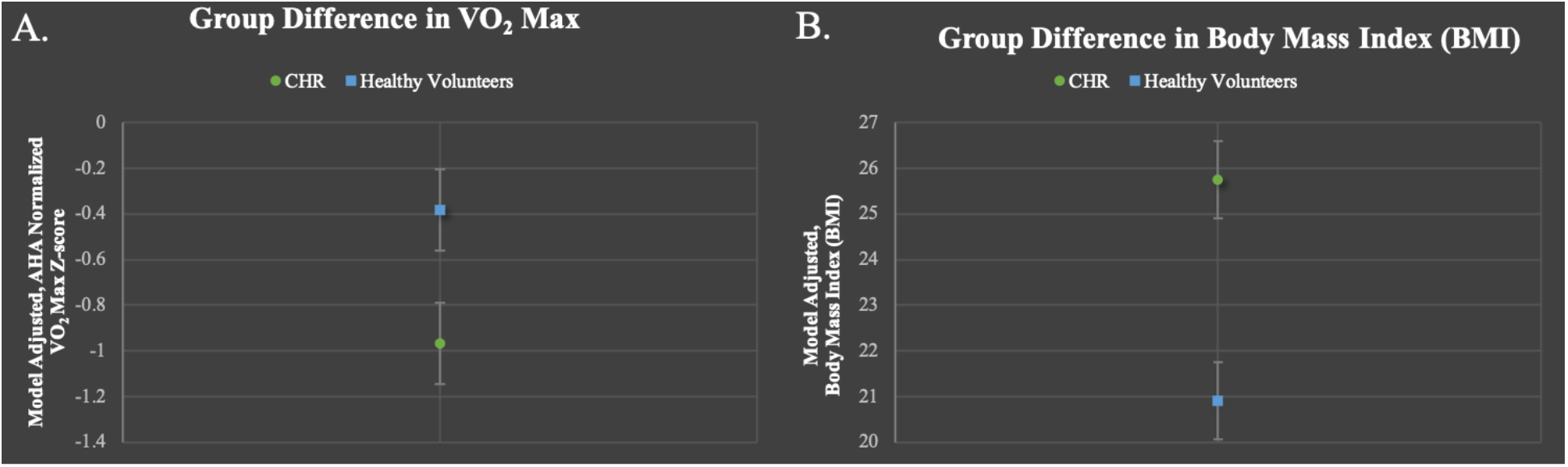
CHR Physical Markers of Fitness and Perceived Fitness Metrics: CHR group differences in VO_2_ max (A) and Body Mass Index (B).

### CHR Self-Reported Fitness Related to Lab-Based Indices of Fitness

In a general linear model, self-reported fitness subscales (Exercise Frequency, Time Spent Exercising, Intensity of Exercise, Perceived Fitness) were entered simultaneously to predict VO_2_max (ml/kg/min) z-scores. Perceived Fitness did not predict VO_2_max values (*p*=.72), but did relate to the intensity of the exercise, *F*(1, 36)=5.17, *p*=.03, *r_partial_*=−.42, *Figure 2a*. There were no other specific exercise features that related to VO_2_max values, *p’s*>.21. In a general linear model, self-reported fitness subscales (Exercise Frequency, Time Spent Exercising, Intensity of Exercise, Perceived Fitness) were entered simultaneously to predict BMI. Perceived Fitness did not predict BMI values (*p*=.47), but did relate to the intensity of the exercise, *F*(1, 36)=5.79, *p*=.02, *r_partial_*=.42, *Figure 2b*, and to the time spent exercising, *F*(1, 36)=4.97, *p*=.03, *r_partial_*=.39, *Figure 2c*. BMI did not relate to the frequency of exercise, *p*=.47.

**Figure 2.**
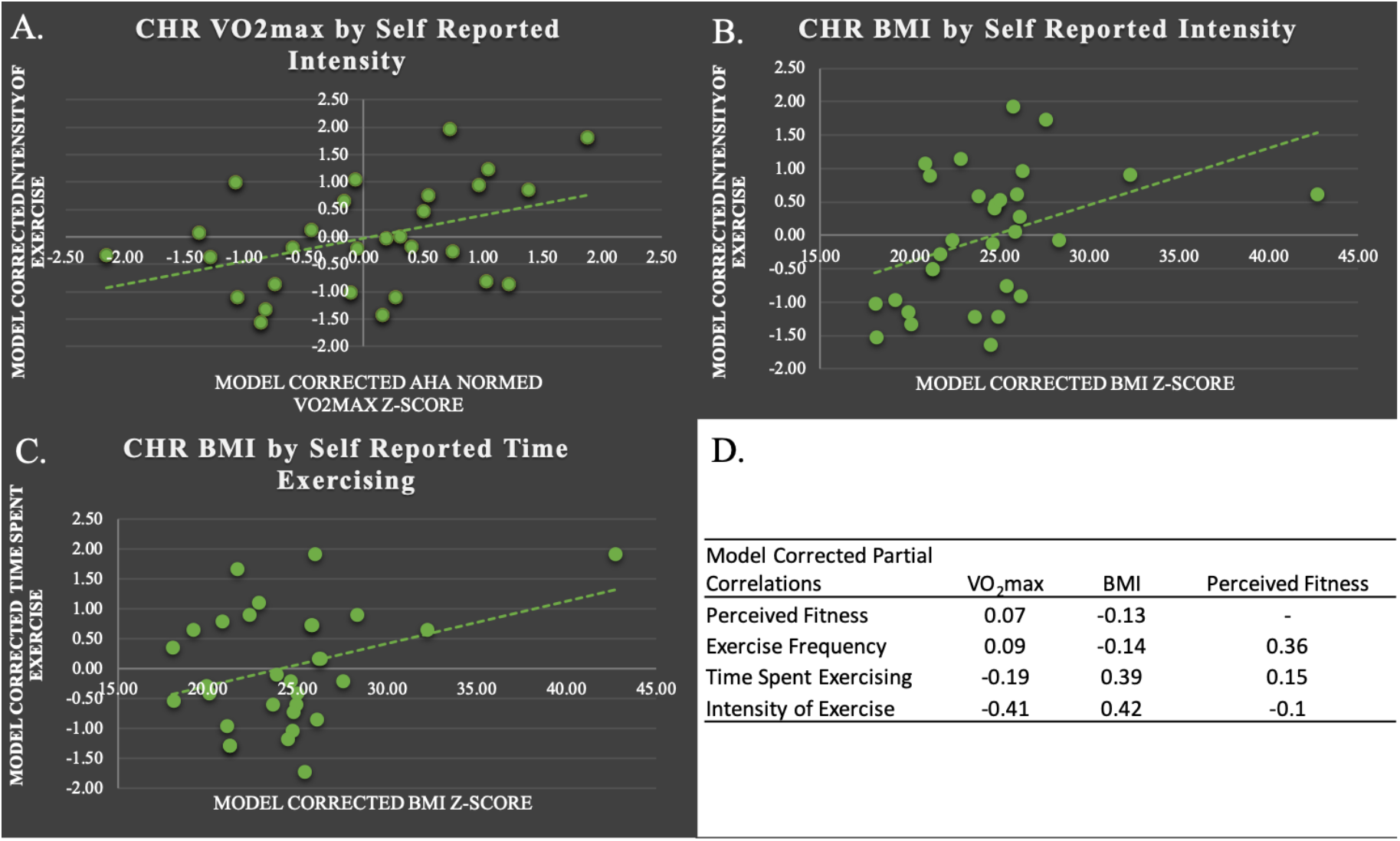
Relationship of Objective Physiological Health to Self-Reported Measures of Fitness: A. Within the CHR group relationshps between VO_2_ max to reported exercise intensity, B. BMI to reported exercise intensity, C. BMI to reported time spent exercising, D. Intercorrelation matrix of the Perceived and Actual Fitness metrics.

### CHR Self-Reported Fitness Related to Symptoms

In a repeated measures general linear model, attenuated psychotic symptoms (positive and negative symptom scales) were related to self-reported fitness subscales (Exercise Frequency, Time Spent Exercising, Intensity of Exercise, Perceived Fitness), accounting for sex. Perceived fitness related to symptoms, even when accounting for specific self-reported fitness behaviors, *F*(1, 34)=5.90, *p*=.02. Additionally, follow-up analyses demonstrated that sex significantly contributed to the overall model, *F*(1, 34)=4.26, *p*=.04, with CHR males subjects showing more severe symptoms (model-corrected mean=11.02, *SEM*=1.05) compared to CHR females (model-corrected mean=7.94, *SEM*=0.84). Age was also examined as a potential contributor to the model, but did not show a significant contribution to the model nor did it change the magnitude or direction of the reported effects, *p*=.78.

### CHR Errors in Perceived Fitness related to Symptoms

In separate t-tests, grandiosity related to errors in perception (how far the actual fitness deviated from the predicted fitness) for both BMI, *t*(38)=2.28, *p*=.04, *Figure 3a*, and VO_2_ max, *t*(38)=2.29, *p*=.04, *Figure 3c*. Avolition related to errors in perception (how far the actual fitness deviated from the predicted fitness) for both BMI, *t*(38)=2.55, *p*=.02,, *Figure 3c*,and VO_2_ max, *t*(38)=2.55, *p*=.02, *Figure 3d*. To examine specificity of these analyses follow-up analyses were conducted examining positive and negative symptom totals; for all analyses it did not relate to errors in perception (how far the actual fitness deviated from the predicted fitness) *p’s*>.12. Follow-up analyses, examined the potential contribution of age and sex to the models above and found that they did not contribute to the model significantly, *p*’s>.51.

**Figure 3.**
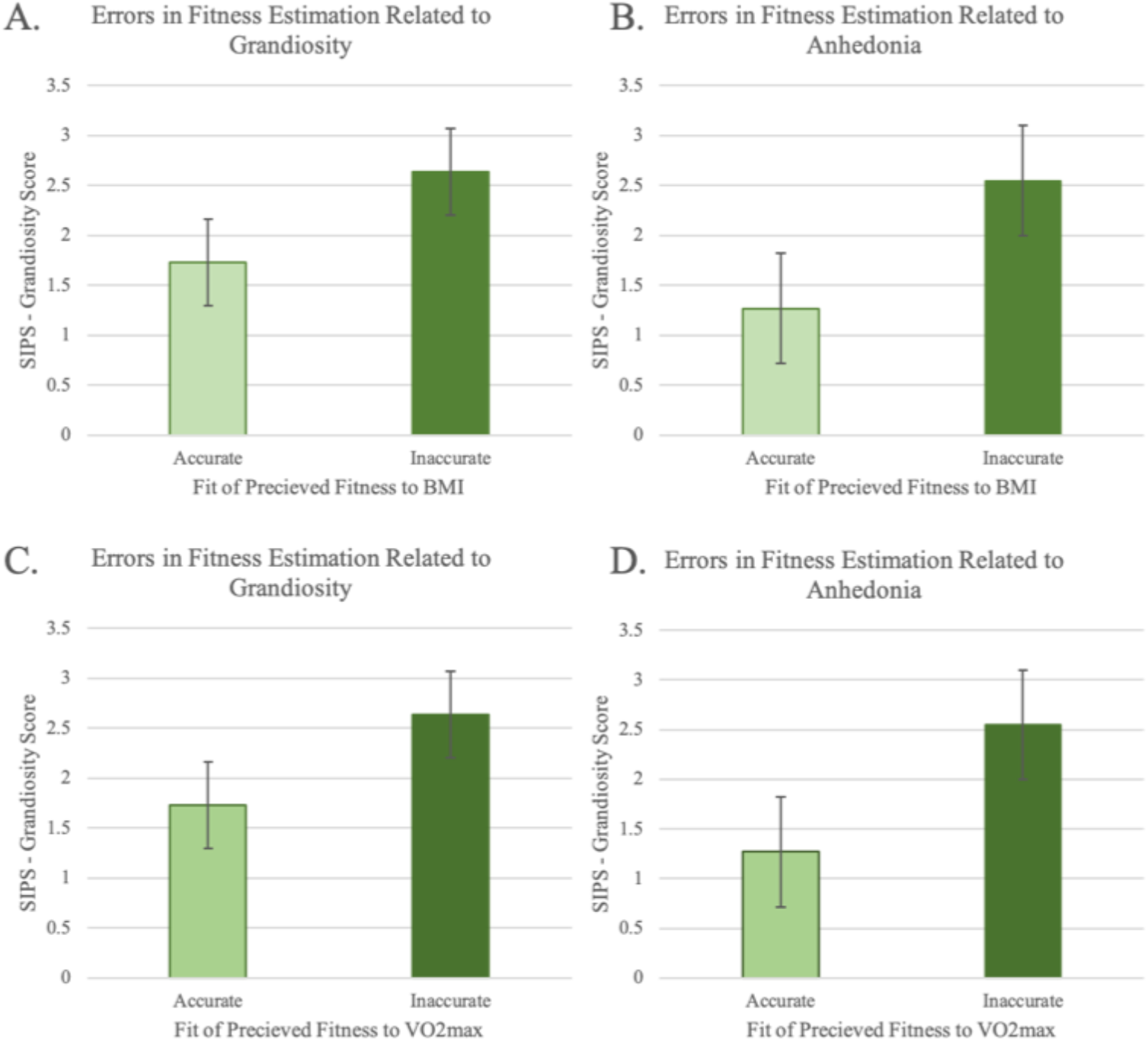
Errors in Perception Related Quartiles Groups (Inaccurate and Accurate) for BMI (top panels) and VO_2_max (bottom panels) to Symptoms of (A,C) Grandiosity and (B,D) Anhedonia

## Discussion

The current paper is the first to take a comprehensive assessment of fitness in CHR individuals. CHR individuals were significantly less fit than peers in terms of lab-based physiological (VO_2_max)^40^ and biometric (BMI) measures of fitness.^41^ Self-reported perceptions of fitness^10,12^ did not reflect either lab-based metric of fitness (VO_2_max, BMI), but specific questions regarding fitness behaviors (i.e., intensity and time spent exercising) did relate to the lab-based metrics of fitness. It is notable, however, that individuals with an elevated BMI^41^ reported higher intensity of exercise,^12^ which may reflect subjective experiences of exercise intensity. This inconsistency further emphasizes the need for more objective accounts of exercise behaviors and fitness.^31^ Finally, distortions in perceived fitness^31^ were related to symptoms related to distortions in perception of self (i.e., grandiosity) and motivation (i.e., anhedonia). In summary, although lab-based measures demonstrate that individuals are less physically fit^40,41^ than peers, the self-report measures of fitness^12^ may reflect a mixture of actual fitness, subjective experience of fitness, and attenuated symptoms.

Results indicated that CHR individuals showed lower levels of fitness on lab based physiological (VO_2_ max)^1,37,40^ and biometric (BMI)^41^ measures in separate analyses, compared to healthy volunteers. This decline in fitness relative to peers emphasizes the potential benefit of exercise as an early, non-invasive intervention in CHR individuals.^6,7,9^ This finding extends the psychosis literature suggesting that a psychosis spectrum diagnosis is associated with lower fitness^1,6,14,19,20^ to include individuals with attenuated psychotic symptoms. Taken together, the emergence of attenuated symptoms co-occurred with a decline in physical fitness may reflect an overall deterioration in neurological fitness or cyclical decline in physiological fitness. This deterioration might lead to a decline in both engagement in fitness activities and neurological health.^1,2,6^ These findings emphasize the importance of lab-based metrics of fitness,^31^ and future studies should evaluate fitness longitudinally to see if these markers of health confer additional risk for conversion and if they may be useful tools in targeting individuals at greatest risk.

Self-report perceived level of fitness^12^ did not reflect lab-based metrics of fitness.^1,37,40,41^ Self-reported intensity of exercise^12^ did reflect physiological fitness as expected; increased self-reported exercise intensity related to an increased VO_2_max capacity.^1,37^ In contrast, self-reported measures of time spent exercising and intensity of exercise^12^ corresponded to an increased BMI, unlike what was expected. This inconsistency with BMI highlights several possibilities. First, this may be a limitation of BMI; BMI does not distinguish between the content of body mass in terms of whether the weight reflects increased muscle mass or body fat.^42–44^ It is possible that individuals with high BMI reflect a grouping of both those individuals with high body fat percentage and elevated weight due to muscle mass.^42,44^ Future studies should consider including estimating body fat percentage, as well as BMI to account for this possibility.^42,44,45^ Alternatively, self-reported levels of intensity and time spent exercising may reflect the subjective experience of the individual.^31,39,46^ As a result, individuals with a higher BMI may experience more difficulty exercising and report objectively less intense exercise as being more intense, and as lasting a longer period of time. In addition to biases of subjective experience, self-report of fitness behaviors may be further distorted by the presence of attenuated symptoms.^31,46^

Self-reports of perceived fitness^12^ reflected overall severity of positive and negative symptoms, even when accounting for other features of health behavior (e.g., frequency and intensity of exercise). This finding suggests that self-reported perception of fitness may, in fact, reflect overall severity of attenuated symptoms of psychosis more than actual fitness.^31^ In exploratory analyses that modeled the difference in perceived fitness to actual fitness, the errors of perceived fitness were related to specific symptoms of self-perception (i.e., grandiosity) and motivation (i.e., avolition). Individuals who were high on symptoms of grandiosity and avolition had higher discrepancy between their self-reported perceived fitness and their predicted perceived fitness scores based on lab-based indices of fitness. Symptoms of grandiosity and impaired motivation distort self-reported assessment of perceived fitness, building on a growing literature indicating clinical influence on some self-report measures.^31,47–50^ Future studies should also examine the possibility that these errors may be effected by deficits in memory and cognitive biases based in mood.^51^

Despite the many strengths and novelty of this study, there are some relevant limitations. Although BMI is a useful metric in distinguishing CHR from healthy controls, it was not entirely clear if increased BMI was reflecting increased muscle mass or body fat percentage.^42–45^ Future studies on this topic should consider including additional metrics of fitness such as waist to hip ratio^20^ or skin fold thickness^45^ to estimate body fat percentage in addition to body mass index. It is also notable that the two contributing sites varied in the specific exercise approach (i.e., treadmill and cycling), which were separately normed with the appropriate population standard.^40,52^ However, this concerned is somewhat mitigated by the equivalent peak heart rates across exercise types, which suggests that the exercises were of roughly equivalent rigor. Further research would benefit from having consistency in their approach to exercise interventions, but this study also serves to demonstrate the importance of population norms^40^ to assist in large multi-study collaborations. Additionally, the sites varied according to age; although the model accounted for variability related to age, it remains possible that age impacts the current findings. It is notable that older age should be related to reduced, not increased, indices of fitness, and so it is possible that the effect is larger than what is reported in the current paper. Finally, the current study was of a similar size to comparable extant literature;^1,9,10,25^ however, the field would benefit from an increased sample size to examine additional variables that may affect fitness, e.g. sex and ethnicity. Additionally, this smaller sample size restricted out sensitivity to symptoms within the CHR group; this led to the restriction of analyses to strong candidate subscales. It is notable that the presence of significant relationships despite this reduced power holds promise for future studies following this line of inquiry.

In conclusion, lab-based metrics of fitness suggest that CHR individuals are less physiologically fit than peers. While this emphasizes the potential of exercise to be a beneficial early intervention, future studies should use caution in depending on self-report measures of fitness. The current study suggests that self-reported measures of fitness may reflect overall severity of symptoms, and that the errors in perceived fitness may, in fact, reflect grandiosity and anhedonia. As a result, future studies of exercise interventions should emphasize objective lab-based metrics (VO_2_max) or concrete features of exercise behavior (time and intensity of exercise) rather than insight-based questions of fitness (perceived fitness). These conclusions may be used to guide the establishment of future exercise interventions: goals of interventions in CHR would be better guided by objective measures of fitness (e.g., goal heart rate and confirmed durations) rather than relying on subject recall^31^ or subjective intensity.^46^

## Acknowledgments

This work was supported by the National Institutes of Mental Health (Grant R01s. MH094650, MH112545–01, MH103231, MH112545, MH094650, R21/R33MH103231). We have no conflicts to disclose.

## References

1. Kimhy D, Vakhrusheva J, Bartels MN, et al. Aerobic fitness and body mass index in individuals with schizophrenia: Implications for neurocognition and daily functioning. Psychiatry Research. 2014;220(3):784–791. doi:10.1016/j.psychres.2014.08.052

2. Kimhy D, Vakhrusheva J, Bartels MN, et al. The Impact of Aerobic Exercise on Brain-Derived Neurotrophic Factor and Neurocognition in Individuals With Schizophrenia: A Single-Blind, Randomized Clinical Trial. Schizophr Bull. 2015;41(4):859–868. doi:10.1093/schbul/sbv022

3. Armstrong HF, Bartels MN, Paslavski O, et al. The impact of aerobic exercise training on cardiopulmonary functioning in individuals with schizophrenia. Schizophrenia Research. 2016;173(1):116–117. doi:10.1016/j.schres.2016.03.009

4. Vakhrusheva J, Marino B, Stroup TS, Kimhy D. Aerobic Exercise in People with Schizophrenia: Neural and Neurocognitive Benefits. Curr Behav Neurosci Rep. 2016;3(2):165–175. doi:10.1007/s40473-016-0077-2

5. Ospina LH, Wall M, Jarskog LF, et al. Improving Cognition via Exercise (ICE): Study Protocol for a Multi-Site, Parallel-Group, Single-Blind, Randomized Clinical Trial Examining the Efficacy of Aerobic Exercise to Improve Neurocognition, Daily Functioning, and Biomarkers of Cognitive Change in Individuals with Schizophrenia. J Psychiatr Brain Sci. 2019;4. doi:10.20900/jpbs.20190020

6. Mittal VA, Vargas T, Juston Osborne K, et al. Exercise Treatments for Psychosis: a Review. Curr Treat Options Psych. 2017;4(2):152–166. doi:10.1007/s40501-017-0112-2

7. Dean DJ, Bryan AD, Newberry R, Gupta T, Carol E, Mittal VA. A Supervised Exercise Intervention for Youth at Risk for Psychosis: An Open-Label Pilot Study. J Clin Psychiatry. 2017;78(9):e1167–e1173. doi:10.4088/JCP.16m11365

8. Scheewe TW, Takken T, Kahn RS, Cahn W, Backx FJG. Effects of Exercise Therapy on Cardiorespiratory Fitness in Patients with Schizophrenia. Medicine & Science in Sports & Exercise. 2012;44(10):1834–1842. doi:10.1249/MSS.0b013e318258e120

9. Mittal VA, Gupta T, Orr JM, et al. Physical activity level and medial temporal health in youth at ultra high-risk for psychosis. Journal of Abnormal Psychology. 2013;122(4):1101–1110. doi:10.1037/a0034085

10. Newberry RE, Dean DJ, Sayyah MD, Mittal VA. What prevents youth at clinical high risk for psychosis from engaging in physical activity? An examination of the barriers to physical activity. Schizophrenia Research. 2018;201:400–405. doi:10.1016/j.schres.2018.06.011

11. Mittal V, Dean D, Gupta T, Bryan A. Aerobic Exercise Intervention for Clinical High-Risk Youth Improves Cognitive and Hippocampal Abnormalities. Schizophr Bull. 2017;43(Suppl 1):S168. doi:10.1093/schbul/sbx024.019

12. Deighton S, Addington J. Exercise practices in individuals at clinical high risk of developing psychosis. Early Intervention in Psychiatry. 2015;9(4):284–291. doi:10.1111/eip.12107

13. Hodgekins, J, French, P, Birchwood, M, et al. Comparing time use in individuals at different stages of psychosis and a non-clinical comparison group | Elsevier Enhanced Reader. doi:10.1016/j.schres.2014.12.011

14. Koivukangas J, Tammelin T, Kaakinen M, et al. Physical activity and fitness in adolescents at risk for psychosis within the Northern Finland 1986 Birth Cohort. Schizophrenia Research. 2010;116(2):152–158. doi:10.1016/j.schres.2009.10.022

15. Phillips LK, Voglmaier MM, Deldin PJ. A preliminary study of emotion processing interference in schizophrenia and schizoaffective disorder. Schizophrenia Research. 2007. doi:10.1016/j.schres.2007.04.003

16. Kimhy D, Delespaul P, Ahn H, et al. Concurrent measurement of “real-world” stress and arousal in individuals with psychosis: Assessing the feasibility and validity of a novel methodology. Schizophrenia Bulletin. 2010. doi:10.1093/schbul/sbp028

17. Leavitt VM, Goldberg TE. Episodic Memory in Schizophrenia. Neuropsychol Rev. 2009;19(3):312–323. doi:10.1007/s11065-009-9107-0

18. Valli I, Tognin S, Fusar-Poli P, Mechelli A. Episodic Memory Dysfunction in Individuals at High-Risk of Psychosis: A Systematic Review of Neuropsychological and Neurofunctional Studies. doi:info:doi/10.2174/138161212799316271

19. Vancampfort D, De Hert M, Myin-Germeys I, et al. Lower cardiorespiratory fitness is associated with more time spent sedentary in first episode psychosis: A pilot study. Psychiatry Research. 2017;253:13–17. doi:10.1016/j.psychres.2017.03.027

20. Shah P, Iwata Y, Caravaggio F, et al. Alterations in body mass index and waist-to-hip ratio in never and minimally treated patients with psychosis: A systematic review and meta-analysis. Schizophrenia Research. 2019;208:420–429. doi:10.1016/j.schres.2019.01.005

21. Fusar-Poli P, Howes OD, Allen P, et al. Abnormal Frontostriatal Interactions in People With Prodromal Signs of Psychosis: A Multimodal Imaging Study. Arch Gen Psychiatry. 2010;67(7):683–691. doi:10.1001/archgenpsychiatry.2010.77

22. Carney R, Cotter J, Bradshaw T, Firth J, Yung AR. Cardiometabolic risk factors in young people at ultra-high risk for psychosis: A systematic review and meta-analysis. Schizophrenia Research. 2016;170(2):290–300. doi:10.1016/j.schres.2016.01.010

23. Bernard JA, Mittal VA. Cerebellar-Motor Dysfunction in Schizophrenia and Psychosis-Risk: The Importance of Regional Cerebellar Analysis Approaches. Front Psychiatry. 2014;5. doi:10.3389/fpsyt.2014.00160

24. Hayes LN, Severance EG, Leek JT, et al. Inflammatory Molecular Signature Associated With Infectious Agents in Psychosis. Schizophr Bull. 2014;40(5):963–972. doi:10.1093/schbul/sbu052

25. Amminger GP, Schäfer MR, Papageorgiou K, et al. Long-Chain ω-3 Fatty Acids for Indicated Prevention of Psychotic Disorders: A Randomized, Placebo-Controlled Trial. Arch Gen Psychiatry. 2010;67(2):146–154. doi:10.1001/archgenpsychiatry.2009.192

26. Labad J, Stojanovic-Pérez A, Montalvo I, et al. Stress biomarkers as predictors of transition to psychosis in at-risk mental states: Roles for cortisol, prolactin and albumin. Journal of Psychiatric Research. 2015;60:163–169. doi:10.1016/j.jpsychires.2014.10.011

27. Brokmeier LL, Firth J, Vancampfort D, et al. Does physical activity reduce the risk of psychosis? A systematic review and meta-analysis of prospective studies. Psychiatry Research. 2020;284:112675. doi:10.1016/j.psychres.2019.112675

28. Optenberg SA, Lairson DR, Slater CH, Russell ML. Agreement of self-reported and physiologically estimated fitness status in a symptom-free population. Preventive Medicine. 1984;13(4):349–354. doi:10.1016/0091-7435(84)90026-4

29. Patterson SM, Krantz DS, Montgomery LC, Deuster PA, Hedges SM, Nebel LE. Automated physical activity monitoring: Validation and comparison with physiological and self-report measures. Psychophysiology. 1993;30(3):296–305. doi:10.1111/j.1469-8986.1993.tb03356.x

30. Baranowski T. Methodologic Issues in Self-Report of Health Behavior. Journal of School Health. 1985;55(5):179–182. doi:10.1111/j.1746-1561.1985.tb04115.x

31. Andorko ND, Rakhshan-Rouhakhtar P, Hinkle C, et al. Assessing validity of retrospective recall of physical activity in individuals with psychosis-like experiences. Psychiatry Research. 2019;273:211–217. doi:10.1016/j.psychres.2019.01.029

32. American College of Sports Medicine, Riebe D, Ehrman JK, Liguori G, Magal M. ACSM’s Guidelines for Exercise Testing and Prescription.; 2018.

33. Bergier J, Kapka-Skrzypczak L, Biliński P, Paprzycki P, Wojtyła A. Physical activity of Polish adolescents and young adults according to IPAQ: a population based study. Ann Agric Environ Med. 2012;19(1):109–115.

34. Lucas-Carrasco R. The WHO quality of life (WHOQOL) questionnaire: Spanish development and validation studies. Qual Life Res. 2012;21(1):161–165. doi:10.1007/s11136-011-9926-3

35. Kane I, Lee H, Sereika S, Brar J. Feasibility of pedometers for adults with schizophrenia: pilot study. J Psychiatr Ment Health Nurs. 2012;19(1):8–14. doi:10.1111/j.1365-2850.2011.01747.x

36. Simons J, Capio CM, Adriaenssens P, Delbroek H, Vandenbussche I. Self-concept and physical self-concept in psychiatric children and adolescents. Res Dev Disabil. 2012;33(3):874–881. doi:10.1016/j.ridd.2011.12.012

37. Balke B, Ware RW. The Present Status of Physical Fitness in the Air Force. SCHOOL OF AVIATION MEDICINE RANDOLPH AFB TX; 1959. https://apps.dtic.mil/docs/citations/ADA036235. Accessed February 20, 2020.

38. Hollenberg M, Ngo LH, Turner D, Tager IB. Treadmill Exercise Testing in an Epidemiologic Study of Elderly Subjects. The Journals of Gerontology Series A: Biological Sciences and Medical Sciences. 1998;53A(4):B259–B267. doi:10.1093/gerona/53A.4.B259

39. Borg G. Borg’s Perceived Exertion and Pain Scales. Champaign, IL, US: Human Kinetics; 1998.

40. Fletcher Gerald F., Balady Gary J., Amsterdam Ezra A., et al. Exercise Standards for Testing and Training. Circulation. 2001;104(14):1694–1740. doi:10.1161/hc3901.095960

41. Calculate Your BMI – Standard BMI Calculator. https://www.nhlbi.nih.gov/health/educational/lose_wt/BMI/bmicalc.htm. Accessed February 20, 2020.

42. Deurenberg P, Weststrate JA, Seidell JC. Body mass index as a measure of body fatness: age- and sex-specific prediction formulas. British Journal of Nutrition. 1991;65(2):105–114. doi:10.1079/BJN19910073

43. Deurenberg-Yap M, Schmidt G, van Staveren W, Deurenberg P. The paradox of low body mass index and high body fat percentage among Chinese, Malays and Indians in Singapore. International Journal of Obesity. 2000;24(8):1011–1017. doi:10.1038/sj.ijo.0801353

44. Malina RM, Katzmarzyk PT. Validity of the body mass index as an indicator of the risk and presence of overweight in adolescents. Am J Clin Nutr. 1999;70(1):131S–136S. doi:10.1093/ajcn/70.1.131s

45. Freedman DS, Sherry B. The Validity of BMI as an Indicator of Body Fatness and Risk Among Children. Pediatrics. 2009;124(Supplement 1):S23–S34. doi:10.1542/peds.2008-3586E

46. Lenka K, David P, Karel K, Zdeněk H. Relationship between subjectively perceived exertion and objective loading in trained athletes and non-athletes. 2015:8.

47. Brent BK, Seidman LJ, Thermenos HW, Holt DJ, Keshavan MS. Self-disturbances as a possible premorbid indicator of schizophrenia risk: A neurodevelopmental perspective. Schizophrenia Research. 2014;152(1):73–80. doi:10.1016/j.schres.2013.07.038

48. Brent BK, Seidman LJ, Coombs G, Keshavan MS, Moran JM, Holt DJ. Neural responses during social reflection in relatives of schizophrenia patients: Relationship to subclinical delusions. Schizophrenia Research. 2014;157(1):292–298. doi:10.1016/j.schres.2014.05.033

49. Damme KSF, Pelletier-Baldelli A, Cowan HR, Orr JM, Mittal VA. Distinct and opposite profiles of connectivity during self-reference task and rest in youth at clinical high risk for psychosis. Hum Brain Mapp. 2019;40(11):3254–3264. doi:10.1002/hbm.24595

50. Cowan HR, McAdams DP, Mittal VA. Core beliefs in healthy youth and youth at ultra high-risk for psychosis: Dimensionality and links to depression, anxiety, and attenuated psychotic symptoms. Dev Psychopathol. 2019;31(1):379–392. doi:10.1017/S0954579417001912

51. Depressed mood in individuals with schizophrenia: a comparison of retrospective and real-time measures. https://www.ncbi.nlm.nih.gov/pmc/articles/PMC4430399/. Accessed March 3, 2020.

52. Loftin M, Sothern M, Warren B, Udall J. Comparison of VO2 Peak during Treadmill and Cycle Ergometry in Severely Overweight Youth. J Sports Sci Med. 2004;3(4):554–560.

